# Unravelling Plankton Adaptation in Global Oceans through the Analysis of Lipidomes

**DOI:** 10.1101/2024.07.22.604538

**Authors:** Weimin Liu, Henry C. Holm, Julius S. Lipp, Helen F. Fredricks, Benjamin A. S. Van Mooy, Kai-Uwe Hinrichs

## Abstract

A recent global survey of planktonic lipids showed a fundamental temperature-mediated regulation of lipid unsaturation in the global oceans [Holm H, et al. (2022) Science 376:1487–1491]. We expand the analysis of this dataset, both spatially and methodologically, to examine diverse environmental stressors across the ocean. Utilizing weighted correlation network analysis, we analyzed 3,164 lipid features in the dataset comprising 930 samples of suspended particulate matter, taken across a broad range of oceanographic conditions and water depths up to 600 meters. A total of 16 lipid clusters being co-expressed across diverse environments were identified. This analysis reveals universal relationships between environmental factors and the lipidome of the planktonic community. The largest lipid cluster, comprising 481 lipid features, including glycerolipids with polyunsaturated fatty acids, exhibited a significant enrichment in polar oceans, suggesting the highest lipid diversity in these ocean regions. Remarkably, marine plankton in these regions employ both desaturation and chain shortening for cold acclimation. Additionally, one lipid cluster strongly linked to the plankton residing in the surface of tropical and subtropical oceans was enriched with non-phosphorus lipids. We suggest this adaptive response enables the plankton to cope with phosphorous scarcity and heat stress. Notably, in the subsurface of these regions, a co-expressed cluster of highly unsaturated lipids is consistent with an enhanced production of polyunsaturated fatty acids by phytoplankton, possibly for low light adaptation. This adaptation is important as it may represent a source of essential fatty acids below the warm sea surface where such vital compounds may be diminished in the warmer future.

**Significance:** Marine plankton is vital for marine ecosystems and climate regulation. We analyzed a large lipidomics dataset of 930 samples collected from global oceans. This allowed us to explore how plankton adapt their lipidomes across different environments. Our findings show distinct lipid clusters correlating with specific environmental conditions, revealing mechanisms like chain shortening to cope with cold stress, enrichment of non-phosphorus lipids in tropical surface waters, and increased polyunsaturated fatty acids in low-light tropical subsurface areas. These adaptations are crucial for understanding how climate change will impact marine ecosystems.

## Introduction

Lipids are essential components of cellular membranes in marine organisms, playing a crucial role in maintaining membrane fluidity, mediating nutrient uptake, and adapting to environmental stressors (1). Lipids, particularly intact polar lipids (IPLs), serve as valuable biomarkers for characterizing and identifying microbial groups in marine ecosystems, enabling the reconstruction of microbial community structure and function (2,3). Understanding the distribution and dynamics of lipids in marine environments is crucial for unravelling ecological dynamics as well as biogeochemical processes occurring in global oceans.

The upper ocean is rich in diverse classes of lipids, associated with various biological functions. Due to their vital role as the interface between cells and their surrounding environments, lipids have received notable attention as indicators of microbial community dynamics in studies of the marine water column and sediments. However, lipids often resist simple classification into taxonomically defined sources or environmentally driven adaptations (4). Certain lipids carry taxonomic information: for example, ornithine-containing lipids are widespread among Bacteria but are absent in Archaea and Eukarya (5). Nonetheless, various studies have revealed that the polar head groups and the core structures of diverse glycerolipids are subject to modifications, enabling organisms to better adapt to changing environmental conditions (6). For instance, living organisms can modulate the degree of unsaturation of acyl groups in response to temperature changes, a phenomenon known as homeoviscous adaptation. A recent study by Holm et al. (7) investigated the average unsaturation of lipid species from ten glycerolipid classes across global oceans, revealing that fatty acid unsaturation is fundamentally linked to temperature.

The interplay between environmental factors and the collective lipidomic response is reminiscent of the regulation of genes caused by environmental forcing. In the case of cold stress, for instance, the substitution of saturated fatty acids with unsaturated ones can be analogously described as the induction of elevated ‘expression’ levels of unsaturated lipids in response to cold stress. Such lipidomic responses can be interpreted as a state of “upregulated expression” of functionally related lipid species, an outcome determined by gene expression regulated by environmental factors, and in the case of cold stress, usually a desaturase (8). Weighted correlation network analysis (WGCNA) has been extensively employed in genomics to group functionally related genes into modules and establish connections with traits and environmental factors (9). WGCNA can also be extended to lipidomics to functionally annotate lipids, explore responses to known stressors, and reveal other unforeseen relationships between lipids and environmental factors. In contrast to networks generated from the similarity of molecular structures based on MS^2^ spectra, such as the Global Natural Products Social Molecular Networking (GNPS) (10,11), which is a frequently used untargeted approach in environmental lipidomics (12,13), WGCNA can establish connections between lipids based on their similarity in “expression” under specific environmental settings. Importantly, the results obtained with WGCNA can provide functional annotations for lipids that seamlessly integrate with the GNPS network, thereby facilitating the annotation of novel lipid biomarkers. This is particularly relevant considering the challenges associated with manual examination or library matching, i.e., the currently employed procedures to annotate lipids in environmental samples due to the lack of appropriate mass spectral databases.

In this study, we apply the weighted correlation network analysis to the extensive dataset published by Holm et al. (7) and Holm and Van Mooy (14). This dataset offers a wealth of information on lipidome profiles across diverse marine environments, encompassing various geographical regions and oceanic conditions. By applying our analytical approach to this comprehensive dataset, we aim to investigate the intricate relationships between lipids and environmental factors on a global scale. Specifically, we seek to elucidate how marine plankton adapt to environmental fluctuations, with a particular emphasis on the influence of temperature variations, nutrient availability, and sunlight exposure.

## Results

The sampled water column in the dataset by Holm and Van Mooy (14), includes the epipelagic zone where the surface mixed layer resides, and the uppermost reaches of the mesopelagic zone. A total of 3,164 lipid features are included in our reanalysis of the mass spectrometry data, including 847 lipid features identified in Holm et al. (7) (designated as annotation type 1; Ann.1), 1,486 additional lipid features (designated Ann.2) identified using LOBSTAHS (15) and LIPIDMAPS (16), and 831 lipid features (designated Ann.3) that do not belong to Ann.1 or Ann.2 but yielded MS^2^ spectra. The analysis of these 3,164 lipid features yielded 16 co-expressed clusters (eigenlipids, referred to as ELs), which are ordered by the number of included lipid features (Table 1). Additionally, a collection of lipid features, labelled as EL0, comprised features that were not associated with any cluster (9).

**Table 1.**
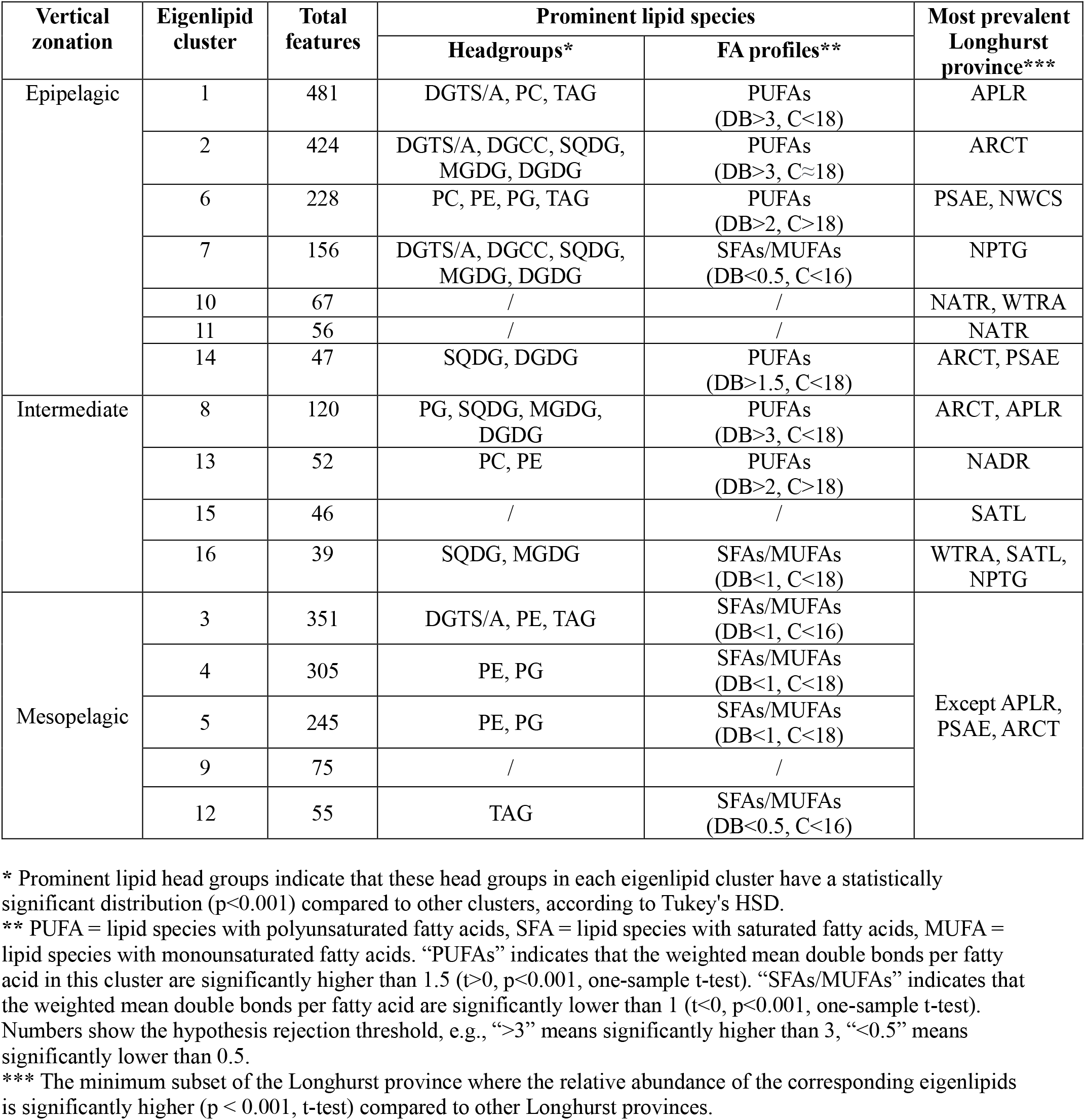
Comprehensive overview of eigenlipid clusters: total features, prominent lipid species, and distribution traits across oceans. See Table S1 for abbreviations of lipid species and Longhurst provinces.

| Vertical zonation | Eigenlipid cluster | Total features | Prominent lipid species |  | Most prevalent Longhurst province*** |
| --- | --- | --- | --- | --- | --- |
|  |  |  | Headgroups* | FA profiles** |  |
| Epipelagic | 1 | 481 | DGTS/A, PC, TAG | PUFAs (DB>3, C<18) | APLR |
|  | 2 | 424 | DGTS/A, DGCC, SQDG, MGDG, DGDG | PUFAs (DB>3, C≈18) | ARCT |
|  | 6 | 228 | PC, PE, PG, TAG | PUFAs (DB>2, C>18) | PSAE, NWCS |
|  | 7 | 156 | DGTS/A, DGCC, SQDG, MGDG, DGDG | SFAs/MUFAs (DB<0.5, C<16) | NPTG |
|  | 10 | 67 | / | / | NATR, WTRA |
|  | 11 | 56 | / | / | NATR |
|  | 14 | 47 | SQDG, DGDG | PUFAs (DB>1.5, C<18) | ARCT, PSAE |
| Intermediate | 8 | 120 | PG, SQDG, MGDG, DGDG | PUFAs (DB>3, C<18) | ARCT, APLR |
|  | 13 | 52 | PC, PE | PUFAs (DB>2, C>18) | NADR |
|  | 15 | 46 | / | / | SATL |
|  | 16 | 39 | SQDG, MGDG | SFAs/MUFAs (DB<1, C<18) | WTRA, SATL, NPTG |
| Mesopelagic | 3 | 351 | DGTS/A, PE, TAG | SFAs/MUFAs (DB<1, C<16) | Except APLR, PSAE, ARCT |
|  | 4 | 305 | PE, PG | SFAs/MUFAs (DB<1, C<18) |  |
|  | 5 | 245 | PE, PG | SFAs/MUFAs (DB<1, C<18) |  |
|  | 9 | 75 | / | / |  |
|  | 12 | 55 | TAG | SFAs/MUFAs (DB<0.5, C<16) |  |

\* Prominent lipid head groups indicate that these head groups in each eigenlipid cluster have a statistically significant distribution ( $p < 0.001$ ) compared to other clusters, according to Tukey's HSD.
\*\* PUFA = lipid species with polyunsaturated fatty acids, SFA = lipid species with saturated fatty acids, MUFA = lipid species with monounsaturated fatty acids. "PUFAs" indicates that the weighted mean double bonds per fatty acid in this cluster are significantly higher than 1.5 ( $t > 0$ , $p < 0.001$ , one-sample t-test). "SFAs/MUFAs" indicates that the weighted mean double bonds per fatty acid are significantly lower than 1 ( $t < 0$ , $p < 0.001$ , one-sample t-test). Numbers show the hypothesis rejection threshold, e.g., ">3" means significantly higher than 3, "<0.5" means significantly lower than 0.5.
\*\*\* The minimum subset of the Longhurst province where the relative abundance of the corresponding eigenlipids is significantly higher ( $p < 0.001$ , t-test) compared to other Longhurst provinces.

EL1 encompasses the highest number of features, totaling 481 (Table 1), while EL6 demonstrates the greatest average abundance across the samples (Figure 1A). Conversely, the relative abundances of certain clusters, such as EL10 and EL15, are nearly negligible, approaching zero in the majority of samples (Figure 1A). Analogous to the concept of “eigengenes,” which represents genes exhibiting similar expression patterns in a given dataset (9), in this analysis “eigenlipids” exhibit comparable distribution patterns across diverse environments. Indeed, these eigenlipids exhibit a remarkable variation in lipid composition (Figure S1 – S3), as well as distinctive vertical (Figure 1B) and geographical distributions (Figure 2) across the global oceans. In terms of their vertical distribution in the upper ocean, these 16 ELs can be broadly categorized as “epipelagic” clusters, “intermediate” clusters, and “mesopelagic” clusters, based on their depth of highest expression (Figure 1B). Regarding their geographical distribution, we analyzed the distribution of these 16 ELs in the context of Longhurst provinces (17) (Table 1, Figure 2).

**Figure 1.**
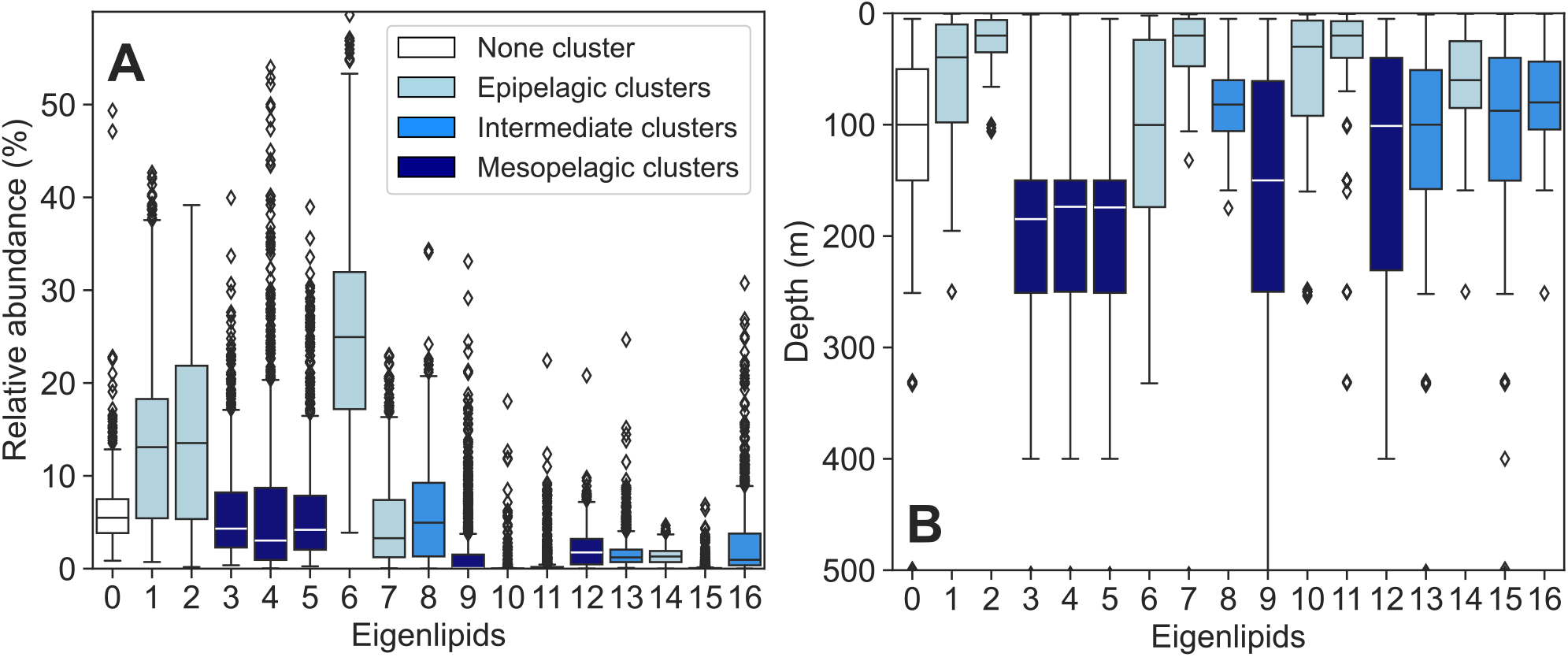
Relative abundances and vertical distribution of eigenlipid clusters in global oceans. (A) Distribution of the relative abundances of eigenlipid clusters across samples. (B) Depth distribution at which each eigenlipid exhibits peak abundance in the water column, showing that these eigenlipids possess depth specificity throughout the upper ocean. For example, EL2 reaches its peak abundance frequently within 50 m below sea surface.

**Figure 2.**
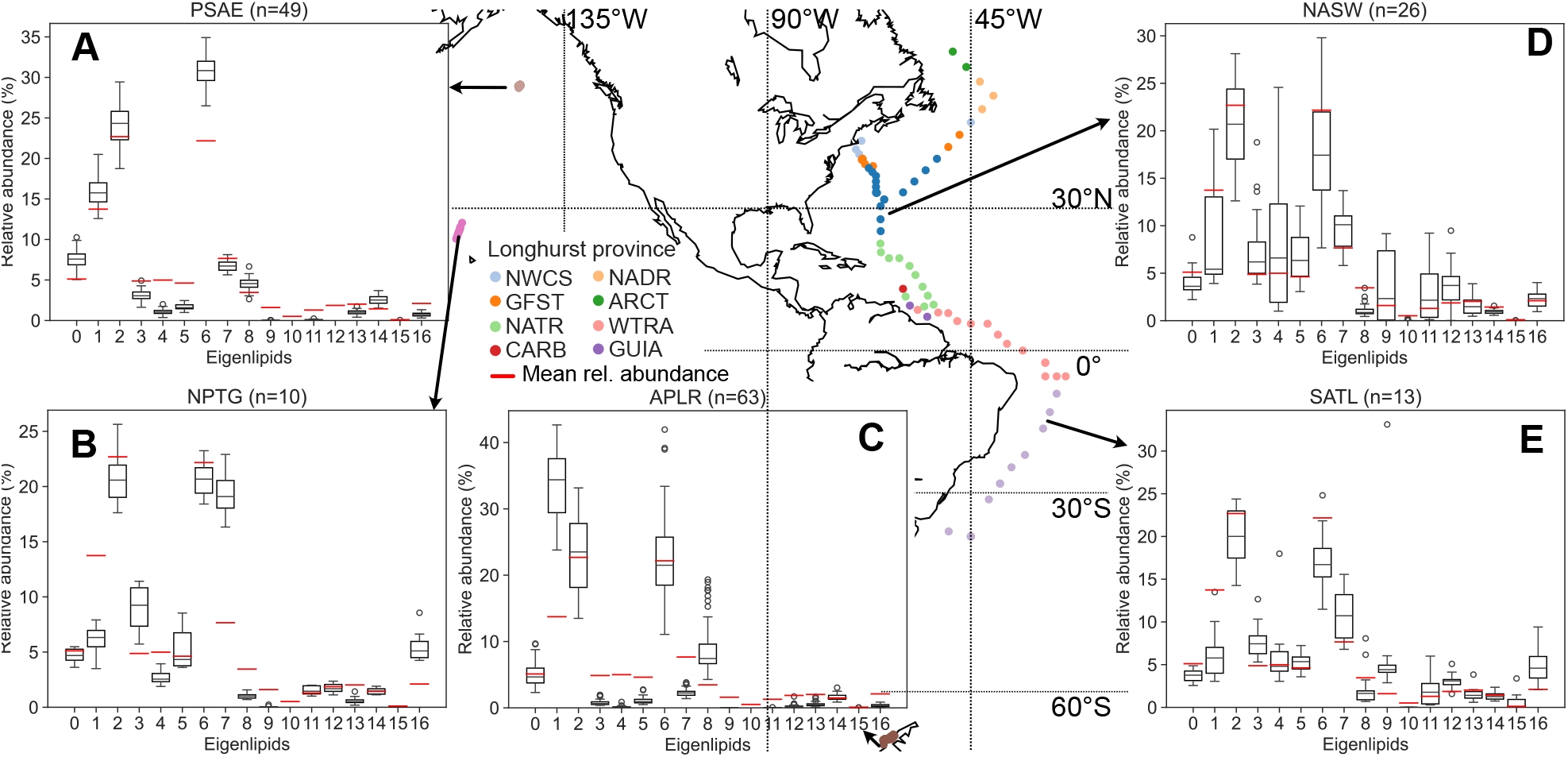
Compositional structure of eigenlipids in the global oceans. The map illustrates dataset sampling locations, with each site color-coded according to the Longhurst province code. Panels A-E depict the distribution of eigenlipid compositions in the mixed layer of specific Longhurst provinces: PSAE (Eastern Pacific Subarctic Gyres), NPTG (North Pacific Tropical Gyre), APLR (Austral Polar), NASW (Northwest Atlantic Subtropical Gyre), and SATL (South Atlantic Gyre), respectively; other Longhurst provinces are defined in the legend and Table S1. The red bar represents the mean relative abundance of eigenlipids across the dataset.

### Epipelagic clusters

Of all epipelagic clusters, EL1, EL2, and EL6 exhibit the highest abundance in polar and subpolar ocean regions (Figure 2A and C). Notably, EL1 emerges as the most prevalent lipid cluster in the Austral polar province, characterized predominantly by triacylglycerols (TAGs) and phosphatidylcholines (PCs) that possess fatty acid chains with an average of fewer than 18 carbon atoms and more than three double bonds per chain (Figure S1 – S2). EL2 and EL6 are the second most prevalent lipid clusters in the Austral polar province, while they dominate in the relatively warmer Eastern Pacific subarctic gyre. The three clusters all contain predominately polyunsaturated fatty acids (PUFA) (Figure S2B). EL6 contain primarily phospholipids and TAGs that possess fatty acid chains with more than 18 carbon atoms and fewer than three double bonds on average. In contrast, EL2 is exclusively dominated by non-phosphorus lipids, including both betaine and glycolipids, containing on average around 18 carbons atoms and more than three double bonds per fatty acid chain.

EL7 is the major epipelagic cluster that exhibits pronounced greater abundance in warm oceanic regions. Its relative abundance remains below 10% in polar and subpolar environments, yet it emerges as one of the most abundant eigenlipid cluster in tropical and subtropical oceans (Figure 2B, D, and E). EL7 is characterized by a predominance of glycolipids containing fully saturated fatty acids (SFA) and/or monounsaturated fatty acids (MUFAs) (Figure S2B). Notably, EL7 exhibits the lowest degrees of unsaturation of all eigenlipid clusters. Furthermore, the glycerolipids in EL7 are dominated by relatively short fatty acid chains, averaging less than 15 carbon atoms per chain (Figure S2A).

EL10 and EL11 appear to be region specific. EL11 shows a relatively higher abundance in the Sargasso Sea compared to other ocean regions (Figure S4). Conversely, EL10 exhibits relatively higher abundance in surface waters near the South American coast (Figure S4). It is noteworthy that none of the glycerolipids identified in the study by Holm et al. (7) are clustered within these clusters. Additionally, the Ann.2 lipids from these clusters are characterized by the absence of phosphorus in their elemental compositions (Figure S3D).

EL14 is ubiquitous in the global oceans, although it only accounts for a small fraction of the lipidome. It contains primarily glycolipids with relatively short fatty acid chains, averaging around 16 carbon atoms per chain.

### Intermediate clusters

While both intermediate clusters EL8 and EL16 are present at sea surface, they frequently exhibit a higher relative abundance in the deep chlorophyll maximum (DCM) zone compared to the sea surface (Figure 1B). EL8 is predominantly composed of glycolipids containing PUFAs with approximately 18 carbons and more than three carbon atoms per chain. Notably, EL8 has the lowest hydrogen-to-carbon ratio among the Ann.2 lipids (Figure S3B), which may result from the inclusion of pigments such as xanthophylls. By contrast, although EL16 is also dominated by glycolipids, it contains primarily SFAs and/or MUFAs with chain lengths fewer than 16 carbons, as opposed to the PUFAs-prevalent EL8. Moreover, EL16 exhibits a higher relative abundance than EL8 in the surface of tropical and subtropical regions (Figure 2B, D, and E).

The other two intermediate clusters EL13 and EL15 are less abundant in the global oceans. EL13 contains primarily phospholipids with relatively long PUFA chains, averaging more than 18 carbon atoms per chain. EL15 contains none of the glycerolipids identified in the study by Holm et al. (7).

### Mesopelagic clusters

EL3, EL4, EL5, EL9, and EL12 are eigenlipids that frequently exhibit significantly higher abundance at the bottom of epipelagic zone and at the top of mesopelagic zone. Their relative abundances combined are almost negligible in the surface of polar and subpolar oceans (Figure 2A and C), while they can reach 30% of the total lipidome in the mixed layer of tropical and subtropical oceans. The glycerolipids included in these eigenlipid clusters contain predominantly SFAs and/or MUFAs (Figure S2B). Notably, glycolipids are almost absent in all mesopelagic clusters. Phospholipids are observed in EL3, EL4, and EL5, with phosphatidylethanolamine (PE) being the major phospholipid in EL3, and both phosphatidylglycerol (PG) and PE being major phospholipids in EL4 and EL5 (Figure S1). Furthermore, alongside the intact polar lipids, EL3 exhibits a substantial abundance of neutral lipids, including ceramides and TAGs (Figure S1). Notably, these neutral lipids contain primarily SFAs and/or MUFAs, resulting in a high H/C ratio (Figure S3B). EL4 are characterized by the predominance of small and polar molecules (Figure S3 A and C). EL12 is primarily composed of TAGs with SFAs and/or MUFAs (Figure S1), while the fatty acid chains in EL12 are longer compared to the those in EL3 (Figure S2A).

## Discussion

The application of weighted correlation network analysis (WGCNA) to the sample set collected by Holm and Van Mooy (14), which includes not only the mixed layer but also extends to the uppermost section of the mesopelagic ocean, has led to the identification of structurally distinctive eigenlipid clusters. These eigenlipid clusters exhibit correlations with environmental data (Figure S5 and S6) that extend beyond the relationship between lipid unsaturation and temperature discussed in Holm et al. (7), indicating a broader range of environmental interactions. Given that there is no *a priori* reason for WGCNA to differentiate between structurally different lipids, the observed structural heterogeneity among eigenlipids suggests that the lipid composition reflect the environmental conditions within the water column. However, it is unclear whether such link arises from microbial adaptation to dynamic environmental conditions or from the presence of distinct microbial communities inhabiting the water column.

Above all, the two abundant eigenlipids EL2 and EL6 appear to have a mixed microbial source. They are characterized by a variety of head groups commonly found in all marine plankton. This includes headgroups such as PE, which are more commonly associated with Bacteria (2,18), as well as PC containing long-chain (C_20_ and C_22_) PUFAs (Table S2), which typically derive from eukaryotic phytoplankton (19). Conversely, the mesopelagic cluster EL3 is dominated by PE with SFA/MUFA chains along with minor amounts of Bacteria-specific ornithine lipids (5), which potentially suggests a primary origin from Bacteria based on published studies on microbial sources of intact polar lipids in marine environments (2,18). Nevertheless, such attribution raises questions about the origins of the co-expressed ceramides and short-chain saturated TAGs with EL3, specifically whether they also primarily originate from Bacteria. Furthermore, both EL8 and EL16 contain primarily canonical chloroplast lipids and exhibit a positive correlation with chlorophyll a (Figure S6E) and divinyl chlorophyll a (Figure S6G), respectively. While chlorophyll a is utilized by a broad spectrum of phytoplankton, divinyl chlorophyll a is predominantly associated with the cyanobacterium *Prochlorococcus* (20). Consequently, it is plausible that EL8 primarily originates from a diverse range of phytoplankton, whereas EL16 may primarily originate from *Prochlorococcus*. Therefore, it appears that eigenlipids may serve not only as markers of microbial adaptations to environmental changes but also, to some extent, as indicators of distinct microbial sources in various marine environments.

### Cold environments favor shorter acyl moieties

Homeoviscous adaption by the desaturation of fatty acid chains in glycerolipids in response to decreasing temperature has been well-documented by Holm et al. (7). Notably, the PUFA-containing EL1 lipids appear to be more responsive to changing temperature than the other two PUFA-containing clusters EL2 and EL6. The abundance ratios of EL1 to EL6 and EL1 to EL2 lipids show that EL1 lipids are less abundant than EL6 and EL2 in tropical and subtropical regions, yet more abundant in polar oceans (Figure S7). Such contrast in lipid abundance despite similar unsaturation may stem from differences in chain length of the fatty acids. Specifically, EL1 lipids have shorter acyl chains than EL2 and EL6 lipids (Figure S2A). In fact, a similar phenomenon is be observed for the predominantly SFA/MUFA-containing epipelagic cluster EL7: the average chain length is shorter in colder regions (Figure S8). Moreover, we find that there is a significant positive correlation between the chain length of all SFA containing membrane lipids and in-situ temperature (Figure 3A), and between the chain length of all PUFA containing membrane lipids and in-situ temperature (Figure 3B). This suggests that, in addition to an increase in unsaturation, colder environments favor shorter acyl moieties. This preference for shorter-chain fatty acids can be attributed to their lower melting transition temperature compared to longer-chain fatty acids (21), allowing them to remain in a fluid state at lower temperatures.

**Figure 3.**
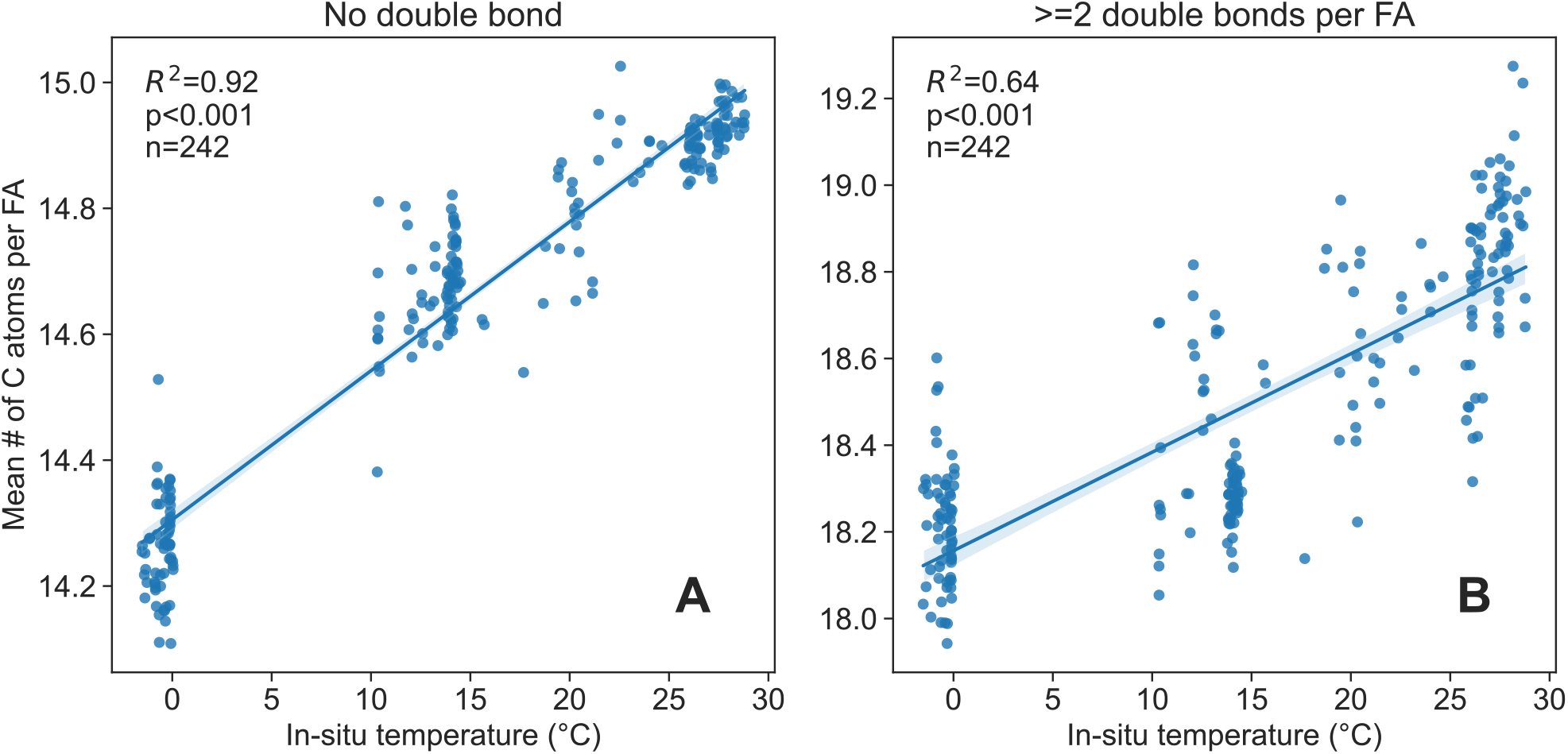
Trends in fatty acid chain length as a function of temperature across all membrane glycerolipids containing fully saturated fatty acids (A) and polyunsaturated fatty acids (B). The weighted mean carbon number per fatty acid (FA) is plotted against temperature for mixed layer samples. The blue line represents linear fits; the inset depicts the coefficient of determination for the linear regression fit using ordinary least squares.

Chain shortening has been reported for eukaryotic phytoplankton. For example, Flaim et al. (22) reported that in a psychrophilic dinoflagellate, TAGs with shorter chain lengths are more abundant at low growth temperatures, while those with longer chain lengths are more abundant under high growth temperature. In fact, a significant difference in the chain length is observed between EL1 and EL6 TAGs, which contain on average 52.7±0.6 and 54.5±1.3 carbon atoms in the combined acyl chains, respectively. Chain shortening has also been observed for polar lipids in certain marine *Synechococcus* strains, which can shorten highly saturated acyl chains. Breton et al. (23) and Pittera et al. (24) reported that these cyanobacteria increase the C_14_/C_16_ fatty acid ratio in glycolipids under low growth temperatures. This observation aligns with the carbon number shift observed in the glycolipids-dominant EL7 cluster from tropical to polar oceans, where the average chain length decreases from 15 to 14.4 carbon atoms per fatty acid (Figure S8). Nonetheless, it remains uncertain whether such difference in chain length indicates that marine plankton are capable of shortening fatty acyl chains as part of their homeoviscous adaption, or if plankton that produce shorter fatty acids are inherently better adapted to cold environments since studies have shown that psychrophiles tend to have shorter acyl chains than mesophiles (25,26). However, regardless of the mechanism responsible for regulating the chain length of unsaturated lipids, it is not limited to the extremely cold polar ocean but is also evident in tropical to subtropical oceans (Figure 3 and S7). Hence, it can be inferred that alterations in lipid chain length may serve as a universal adaptation strategy, alongside lipid unsaturation, for marine plankton as a collective response to temperature fluctuations.

### Accumulation of non-phosphorus lipids in warm waters

Polar oceans are nutrient replete and highly productive. Conversely, tropical and subtropical open oceans generally exhibit stratification, resulting in limited nutrient availability at the surface (27). A pertinent example is the Sargasso Sea, located within the Northern Atlantic subtropical gyre, which is notably depleted of phosphate, measuring at less than 10 nmol/L (28). In response to such phosphate scarcity, plankton have been observed replacing phospholipids with various non-P lipids to reduce their phosphorus requirements (29,30). None of the previously reported substitute lipids have been detected in the lipid cluster unique to the Sargasso Sea, i.e., EL11. Nonetheless, EL7, which predominantly contain non-phosphorus lipids, show relatively high abundance in this region (Figure 2D and S4). Given that EL7 contains lipids reported as substitute lipids utilized by phytoplankton in the Sargasso Sea (29), it is plausible to hypothesize that its increased abundance in this region is attributable, in part, to the substitution of phosphorus-containing lipids with their non-phosphorus counterparts.

Notably, the cluster EL7 is not only abundant in the Sargasso Sea, but also in the other ocean regions. For example, EL7 exhibits great abundance in the Northern Pacific tropical gyre (Figure 2B), where phosphorus is not the limiting nutrient (31). This indicates that the accumulation of non-phosphorus lipids in EL7 is influenced by environmental factors beyond phosphorus availability. In fact, a multivariate linear regression analysis on the rank-transformed data revealed that in-situ temperature had a stronger coefficient (0.88) with EL7 compared to phosphate concentration (0.12), indicating a greater impact of temperature on the relative abundance of EL7 (Figure 4). Our results strongly suggest that the accumulation of EL7 lipids in the global oceans is predominantly influenced by increasing temperature rather than phosphorus limitation.

**Figure 4.**
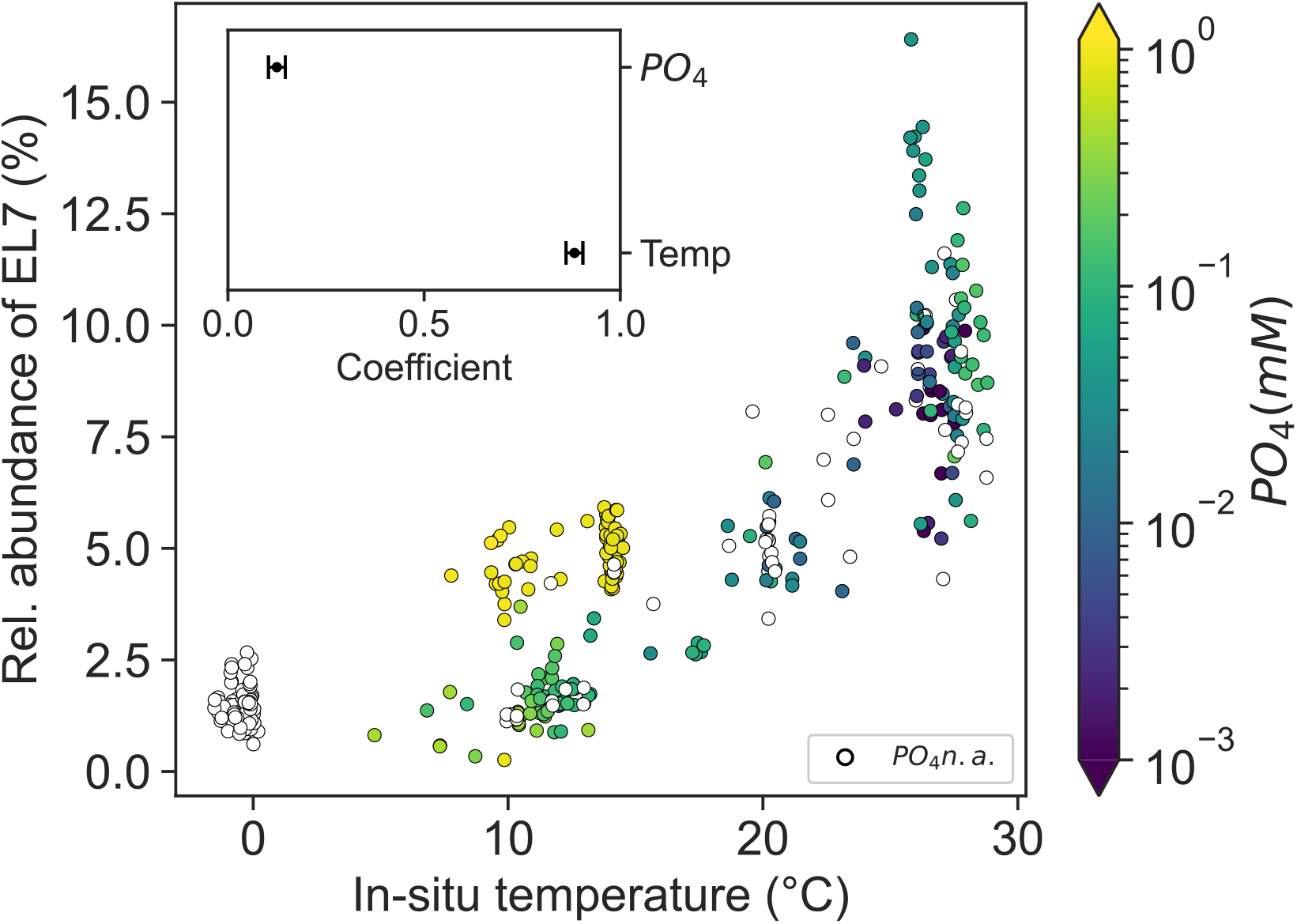
Influence of temperature and phosphate concentration on the relative abundance of the phosphorus-depleted eigenlipid cluster EL7 in global oceans (0-50 m). The color scale represents the concentration of phosphate in the water column, while open circles indicate the unavailability of phosphate concentration data for corresponding samples. The inset shows the coefficients of the multivariate linear regression analysis between the rank-transformed relative abundance of EL7 and rank-transformed environmental variables: in-situ temperature and PO_4_ concentration (adjusted R^2^ = 0.925, p<0.001, n=251).

EL7 is dominated by glycolipids with SFA/MUFA chains. Since most glycolipids are typically found in chloroplast, their presence indicates the accumulation of these non-P lipids could be associated with photosynthesis. Balogi *et al*. (32) refers to saturated MGDG as “heat shock lipid” that accumulates in certain cyanobacteria strains upon exposure to heat and light, preserving the functionality of thylakoid membranes. Indeed, EL7 exhibits a stratified vertical distribution pattern at sea surface (Figure 1B) where light intensity is stronger than the subsurface. Consequently, we propose that the accumulation of these glycolipids at warmer temperatures may result from a collective response by phytoplankton to maintain their photosynthetic capacity. Furthermore, it is important to note that non-P lipids are generally less efficiently exported to the deeper ocean compared to phospholipids (33). Overall, it is plausible that the accumulation of non-P lipids in warm temperature could influence the elemental stoichiometry of the biological pump in a progressively warmer future ocean.

### Deep chlorophyll maximum: A crucial PUFA reservoir in tropical and subtropical oceans

Despite all three clusters containing predominately chloroplast lipids, the intermediate clusters EL8 and EL16 show distinctive vertical distribution patterns compared to EL7, characterized by a higher abundance in the DCM zone of tropical and subtropical oceans (Figure S4). Such elevated abundance could either be attributed to increasing phytoplankton biomass or the photoacclimation of phytoplankton, which increase their cellular chlorophyll content through thylakoid stacking (34,35). Notably, although both EL8 and EL16 show an increase in the DCM zone, the PUFA-containing EL8 cluster shows a steeper increase from the surface to the DCM compared to the SFA/MUFA-containing EL16 cluster (Figure 5). This more pronounced response in EL8 could be linked to its higher PUFA content, as PUFAs have been shown to increase under low-light conditions (36-38). Alternatively, the differing responses between EL8 and EL16 may be attributed to the specific association between EL16 and divinyl chlorophyll-a (Figure S6G). It is possible that the photosystems associated with divinyl chlorophyll may differ in their adaptation to low-light conditions compared to those associated with monovinyl chlorophyll. Specifically, the photosystems utilizing divinyl chlorophyll, which more efficiently harvests blue light (39), might confer less adaptation stress compared to those utilizing monovinyl chlorophyll.

**Figure 5.**
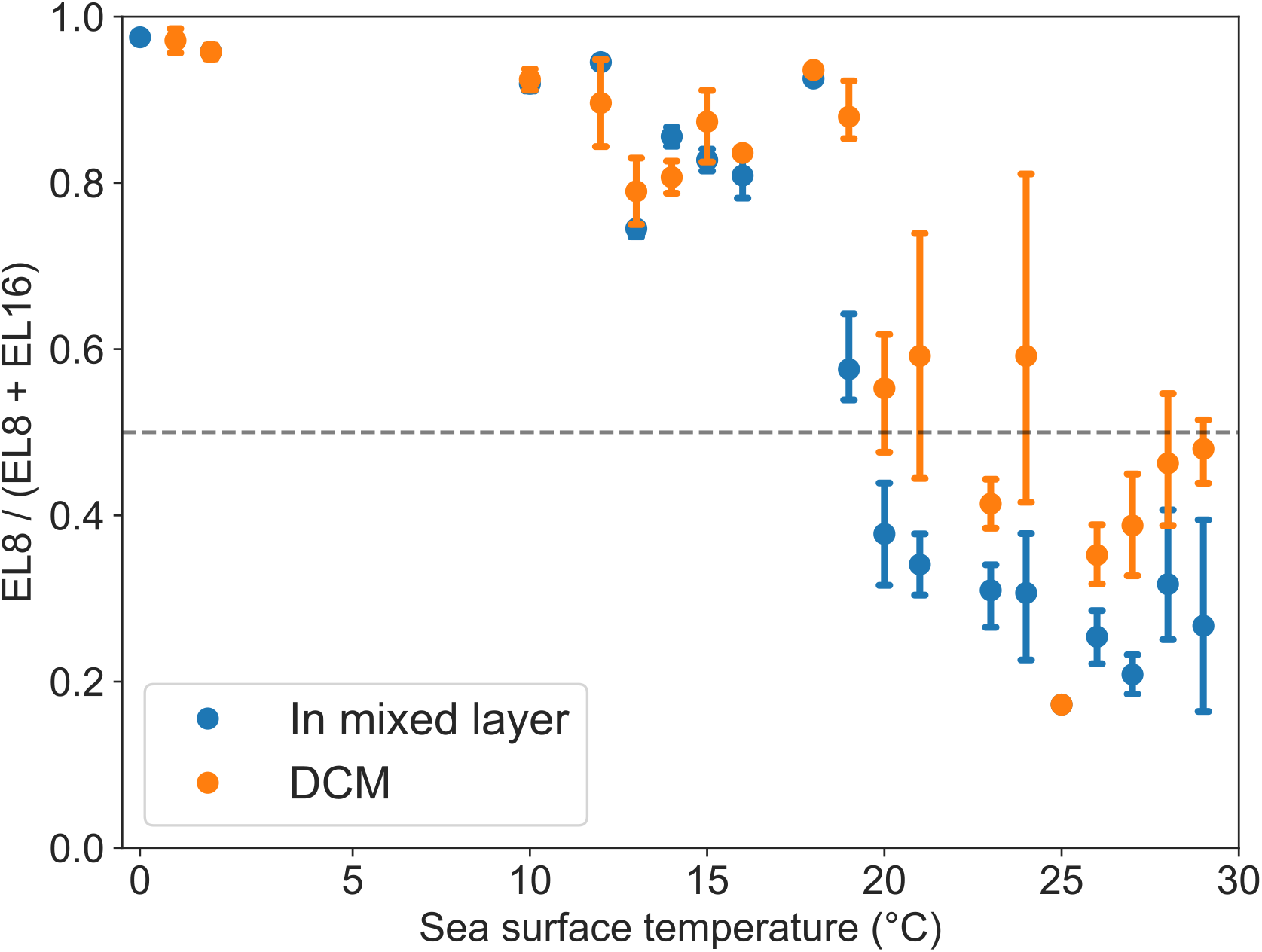
Shifts in the predominance of eigenlipid clusters, EL8 and EL16, from cold to tropical sea surface temperatures (SST) across the global oceans within the mixed layer and the deep chlorophyll maximum (DCM). The abundance of eigenlipid EL8, characterized by its high polyunsaturated fatty acids (PUFAs) content, is plotted against eigenlipid EL16 on the y-axis, represented as EL8/(EL8 + EL16). Error bars indicate the 95% confidence interval of the estimated mean ratio at respective sea surface temperatures.

Nevertheless, such increased PUFA content in the DCM zone suggest that this zone could be an important PUFA reservoir in a warmer ocean. In fact, the concentration of lipids containing eicosapentaenoic acid calculated in Holm et al. (7) indicates that more frequently, their concentrations peak in the subsurface near the depth of peak chlorophyll-a abundance, rather than in the mixed layer of tropical and subtropical oceans (Figure S9). These findings underscore the necessity for ongoing research to elucidate the mechanisms governing PUFAs production and fate, as well as how the penetration of anthropogenically produced heat to these depths influences PUFA production. Such knowledge will be critical for predicting and managing future shifts in marine lipid distributions and their ecological impacts. Nonetheless, the enhanced PUFA production below warmer surface waters, possibly resulting from low-light adaptations, could potentially in part mitigate the loss of essential fatty acids in the anticipated warmer future as described in Holm et al. (7).

## Materials and Methods

### Samples and instrumental analysis

In this study, we used the mass spectrometry data (14) published along with Holm et al. (7) where the procedures for sample collection, extraction, and instrumental analysis have been described in detail.

### Data treatment

The entire process of data manipulation, including both the intermediate and final datasets, is publicly accessible through Zenodo (40). The repository serves as a comprehensive source of the data in its processed form, ensuring transparency and reproducibility of the analysis.

### Feature extraction

In accordance with the initial data processing steps outlined in Holm et al. (7), we employed *xcms* (41) with identical parameters as those specified in the publication. This included utilizing *xcms* for feature extraction, including peak grouping, retention time correction, and peak alignment. Additionally, we collected MS^2^ spectra from the measurements using *xcms* with the *maxTIC* option for the purpose of integrating our functional annotation with GNPS.

### Blank subtraction

To mitigate the presence of potential contaminant signals, we implemented a procedure to remove them. Specifically, we subtracted the peak intensities of features in samples from those in blank samples. Features that exhibited median peak intensities less than 20-fold higher in the samples compared to the blank samples were subsequently eliminated from the analysis.

### Feature grouping and annotation

Consistent with the feature grouping step in Holm et al. (7), the R package CAMERA (42) was employed in this study to annotate potential adducts, isotopologues, and in-source fragments. Feature annotation in this study include three parts with varying quality:

Ann. 1. The robust annotation provided in Holm et al. (7) that includes various glycerolipids.

Ann. 2. Annotations of case codes C2a and C2b (multiple adducts were observed for formulae) generated using a modified version of LOBSTAHS (15), which included an extension of the library to cover ether-linked lipids, along with additional lipids curated in LIPIDMAPS (16). However, it should be noted that the adduct hierarchy for the lipids in the extended libraries cannot be comprehensively validated. Consequently, annotations made using the extended library are, at best, accurate only up to the chemical formula level.

Ann. 3. Annotation generated using custom in-house annotation scripts that rely on MS^2^ spectrum similarity. This involved comparing the MS^2^ spectra obtained from *xcms* against a curated MS/MS library, which was sourced from MS-DIAL (43) and employed the similarity measure *CosineGreedy* as described in Huber et al. (44). In addition, we utilized GNPS (10) to construct a molecular network based on structural similarity.

It is imperative to acknowledge that, due to the varying confidence levels associated with distinct annotation methods, the discussion pertained to specific lipid clusters predominantly revolved around Ann. 1 lipids, with the elemental compositions being a combination of Ann.1 and Ann. 2 lipids. It is worth noting that Ann. 3 lipids are solely included as a confirmation of molecular structure when deemed necessary and as a means to establish a link between the co-expression network and the molecular network generated using GNPS. It is important to clarify that our primary objective in this study is not to achieve precise annotations, as the co-expression network analysis does not necessitate such precision. Instead, the feature annotation step serves as a reference. Moreover, while the annotations may not cover all compounds, we consider this a necessary compromise between including a wide range of lipid signals and eliminating any potential contaminant signals that might persist even after blank subtraction.

### Weighted correlation network analysis

In the weighted correlation network analysis, we utilized the relative abundances of the features. Before conducting the correlation network analysis, we applied a center log ratio transformation.

We then performed a weighted correlation network analysis (WGCNA) per batch (i.e., samples collected from the Atlantic Oceans, the Antarctic, and the Pacific Ocean were analyzed separately, Text S1). Detailed information regarding the analysis parameters and the quality metrics can be found in the supplementary material (Text S2) and the accompanying repository (40). Moreover, while the eigenlipids were derived from samples collected in the Atlantic Ocean (Figure S4), additional data points from the Antarctic (cruise LMG1810) and North Pacific (cruise KM1709 and RR1813) were added after performing WGCNA.

In addition, a separate analysis that applies WGCNA in the mixed layer was performed to validate the idea of employing lipid co-expression as a tool to uncover the collective planktonic response to varying environmental conditions (Text S3).

## Acknowledgments

We thank everyone involved in creating and curating the dataset utilized in this study. Special thanks to Dr. Lars Wörmer for his valuable comments at the beginning of this research. This study was funded by Deutsche Forschungsgemeinschaft (DFG, German Research Foundation) under Germany’s Excellence Strategy—EXC-2077—390741603.

## Data and Code Availability

All the data relevant to this study, as well as all the code used to generate the data, and the figures presented in the manuscript can be found in our data repository on zenodo [https://doi.org/10.5281/zenodo.12790226].

## Author Contributions

W.L., H.C.H., B.A.S.V.M., and K.H. designed research. W.L., H.C.H., J.S.L., H.F.F., B.A.S.V.M., and K.H. analyzed data. W.L. performed research and wrote the draft manuscript. H.C.H., J.S.L., B.A.S.V.M., and K.H. substantially revised the draft manuscript. All authors read and approved the final version of the manuscript.

