## Supplementary Information for "Unravelling Plankton Adaptation in Global Oceans through the Analysis of Lipidomes"

**Text S1: Batch effects**

The samples analyzed in Holm et al. [Holm H, et al. (2022) Science 376:1487–1491] were collected from diverse cruises and regions, encompassing the Atlantic Ocean, the South Ocean, and the Pacific Ocean. Given the substantial time span over which these samples were measured, the possibility of batch effects cannot be discounted. Furthermore, it's important to note that the original measurements were not designed for untargeted analysis. Consequently, it is challenging to ascertain whether the variances observed among geological batches genuinely reflect biogeochemical signals or are attributable to batch effects, as samples from different regions were analyzed at different times.

To mitigate potential interference from batch effects, we opted to conduct weighted correlation network analysis separately for each geological batch. Our primary focus lies on the Atlantic batch, which boasts the largest sample size and encompasses the most extensive range of geochemical variances. Results obtained from the other batches are also included to complement our primary findings from the Atlantic batch.

**Text S2: WGCNA analysis**

The study utilized the WGCNA R package (version 1.69) for investigating the lipidomics data. The lipidomics data used in this study was generated from the Atlantic region. The data pre-processing step involved loading the data using the *fread* function from the data.table package and converting it into a matrix. Hierarchical clustering was then performed using the *hclust* function from the stats package to assess the presence of outliers. The resulting dendrogram was saved as a scalable vector graphics file for further analysis and reference.

The next step in the analysis involved selecting an appropriate soft-thresholding power to construct the network. A range of soft-thresholding powers (1-20) was tested to identify the optimal value. The *pickSoftThreshold* function from the WGCNA package was utilized for this purpose. The analysis utilized the signed network type and *bicor* correlation function. The resulting network topology fit indices were then visualized using a line plot. A scatter plot was also generated to depict the mean connectivity as a function of the soft-thresholding power. The resulting plot was saved as an SVG file for later reference and analysis.

The constructed network was obtained by applying the *blockwiseModules* function from the WGCNA package. The soft-thresholding power value of 12 was chosen based on the previous analysis. The analysis utilized a *bicor* correlation type and a signed TOMType. Other parameters used for construction included a maximum p value for outlier identification of 0.05 and a minimum module size of 20. The resulting module dendrogram was visualized and saved as an SVG file for further analysis.

The module information, including the assigned colors, was saved as a CSV file using the fwrite function from the data.table package. The resulting CSV file was stored in the project directory. Additionally, the module eigenvalues were also saved as a separate CSV file for further investigation and analysis.

**Text S3: Validating WGCNA: Unravelling the link between lipid unsaturation and temperature in surface mixed layer.**

Our principal component analysis (PCA) biplot, shown in Figure S10, provides a comprehensive visualization of the intricate interplay between eigenlipid distribution and the contextual environmental factors. Notably, PC1, which accounts for 44.26% of the total variance in the oceanic lipidome dataset, emerges as a pivotal determinant. Further investigation of the contributing components to PC1 reveals that temperature is the predominant factor. Hence, we establish a clear relationship between eigenlipid composition and temperature, highlighting the potential impact of this environmental factor on lipidome dynamics. To further comprehend the variations in eigenlipid composition in response to temperature changes, we examine the relationships among the eigenlipids. Our analysis reveals that EL0 exhibits a near-orthogonal relationship with PC1, indicating its relative stability amidst temperature fluctuations. On the other hand, EL1 demonstrates a positive correlation with PC1, suggesting its enhanced prevalence in colder aquatic environments. Conversely, EL2 displays a negative correlation with PC1, thereby attributing its higher abundance to warmer water conditions (Figure S11).

In order to investigate the partitioning of the species annotated by Holm et al. [Holm H, et al. (2022) Science 376:1487–1491] within lipid clusters in this study, we have included Figure S12 as supporting evidence. It is worth noting that, except for GADG, there is an observed decrease in the number of double bonds from eigenlipids that display a negative correlation with temperature to those that show a positive correlation with temperature. This observed pattern is in concordance with the findings of Holm et al. [Holm H, et al. (2022) Science 376:1487–1491], which suggest that glycerolipids exhibiting a decrease in abundance with increasing temperature generally possess a higher total number of double bonds compared to those without a negative correlation with temperature. The agreement between our findings and those of Holm et al. [Holm H, et al. (2022) Science 376:1487–1491] provides validation for the assertion that WGCNA has the ability to discern and elucidate the impact of environmental factors on the co-expression of the lipidome.

*Table S1. List of abbreviations for lipid species and Longhurst provinces used*

| Abbreviations | Definitions | Abbreviations | Definitions |
|---|---|---|---|
| DGTS/A | Diacylglyceryl trimethylhomoserine/diacylglyceryl hydroxymethyl-N,N,N-trimethyl-β-alanine | APLR | Austral polar |
| DGCC | Diacylglyceryl-3-O-carboxyhydroxymethylcholine | ARCT | Atlantic Arctic |
| DGDG | Digalactosyldiacylglycerol | CARB | Caribbean |
| MGDG | Monogalactosyldiacylglycerol | GFST | Gulf Stream |
| PC | Phosphatidylcholine | GUIA | Guianas coast |
| PE | Phosphatidylethanolamine | NADR | North Atlantic Drift |
| PG | Phosphatidylglycerol | NASW | Northwest Atlantic subtropical gyre |
| SQDG | Sulfoquinovosyl diacylglycerol | NPTG | North Pacific Tropical gyre |
| TAG | Triacylglycerol | NATR | North Atlantic tropical gyre |
| | | NWCS | Northwest Atlantic shelves |
| | | WTRA | Western tropical Atlantic |
| | | PSAE | Eastern Pacific subarctic gyre |
| | | SATL | South Atlantic gyre |

*Table S2. Distribution of fatty acid composition in the form of "total number of carbon atoms (25$^{th}$ percentile-75$^{th}$ percentile):total number of double bonds (25$^{th}$ percentile-75$^{th}$ percentile) grouped by glycerolipid classes in each eigenlipid cluster. Note that the values refer to the fatty acid moieties excluding the glycerol moieties.*

| ELs | DGCC | DGDG | DGTS/A | MGDG | PC | PE | PG | SQDG | TAG |
|---|---|---|---|---|---|---|---|---|---|
| 1 | 36-38:8-10 | 38-40:3-9 | 31-37:4-8 | / | 34-38:8-10 | 32-36:8-10 | 39-41:2-3 | 31-37:3-6 | 48-54:6-13 |
| 2 | 36-40:5-9 | 34-36:3-8 | 33-40:3-8 | 34-34:1-9 | 32-33:4-5 | / | / | 32-33:0-5 | / |
| 3 | / | / | 37-43:0-1 | / | 39-40:3-5 | 33-34:1-1 | / | / | 40-47:0-2 |
| 4 | / | / | / | / | / | 28-31:1-2 | 31-35:0-2 | / | / |
| 5 | / | / | / | / | / | 28-32:0-2 | 33-37:0-2 | / | / |
| 6 | / | / | / | / | 37-44:6-10 | 37-41:4-10 | 31-32:1-2 | / | 51-60:5-11 |
| 7 | 34-37:1-2 | 29-30:0-0 | 30-39:0-4 | 30-32:0-1 | / | / | / | 29-31:0-0 | / |
| 8 | / | / | / | 34-36:6-9 | / | 32-36:5-7 | 34-34:4-4 | 34-36:5-9 | / |
| 9 | 29-30:0-2 | / | / | / | / | / | / | / | / |
| 10 | / | / | / | / | / | / | / | / | / |
| 11 | / | / | / | / | / | / | / | / | / |
| 12 | / | / | / | / | / | / | / | / | 53-58:1-3 |
| 13 | / | / | / | / | 32-38:2-4 | 38-40:6-6 | / | / | / |
| 14 | 35-37:5-6 | 32-34:3-7 | / | 34-36:3-4 | / | / | / | 32-33:3-4 | / |
| 15 | / | / | / | / | / | / | / | / | / |
| 16 | / | / | / | 28-30:2-3 | / | / | / | 29-31:1-2 | / |

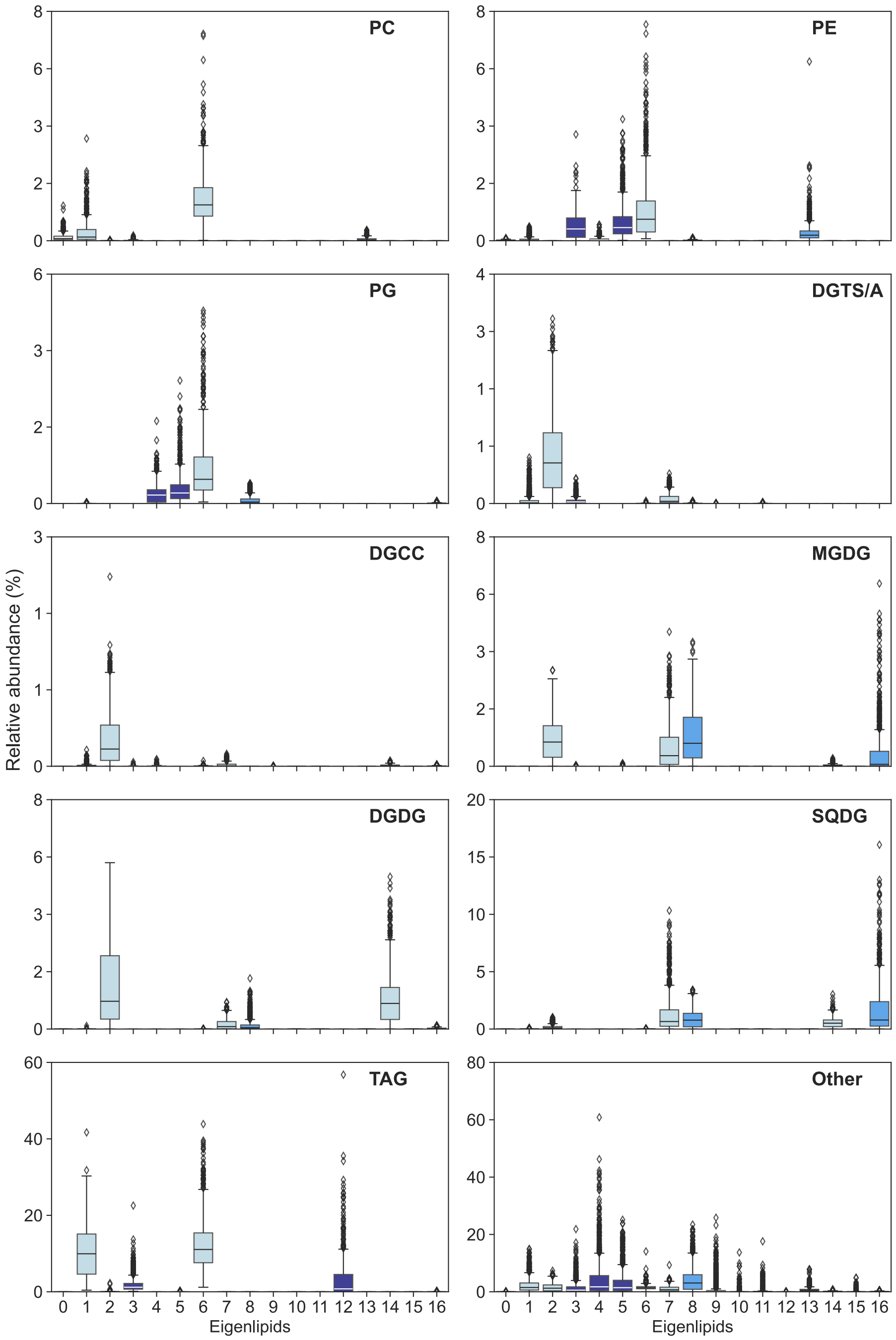


*Figure S1. Distribution of lipid head groups in each eigenlipid identified throughout the sampled water column (i.e., both in and below mixed layer). The y-axis represents the relative abundance of each headgroup in the respective eigenlipid, calculated from the total peak intensities of lipids containing the corresponding headgroup and belonging to the respective eigenlipid cluster, normalized to the total peak intensities of Ann.1, Ann.2, and Ann.3 lipids. Note that, except for 'Other', all relative abundances are corrected against their respective response factors. The x-axis denotes the eigenlipid clusters identified by WGCNA. Note that 'Other' denotes both Ann.2 and Ann.3 lipids.*

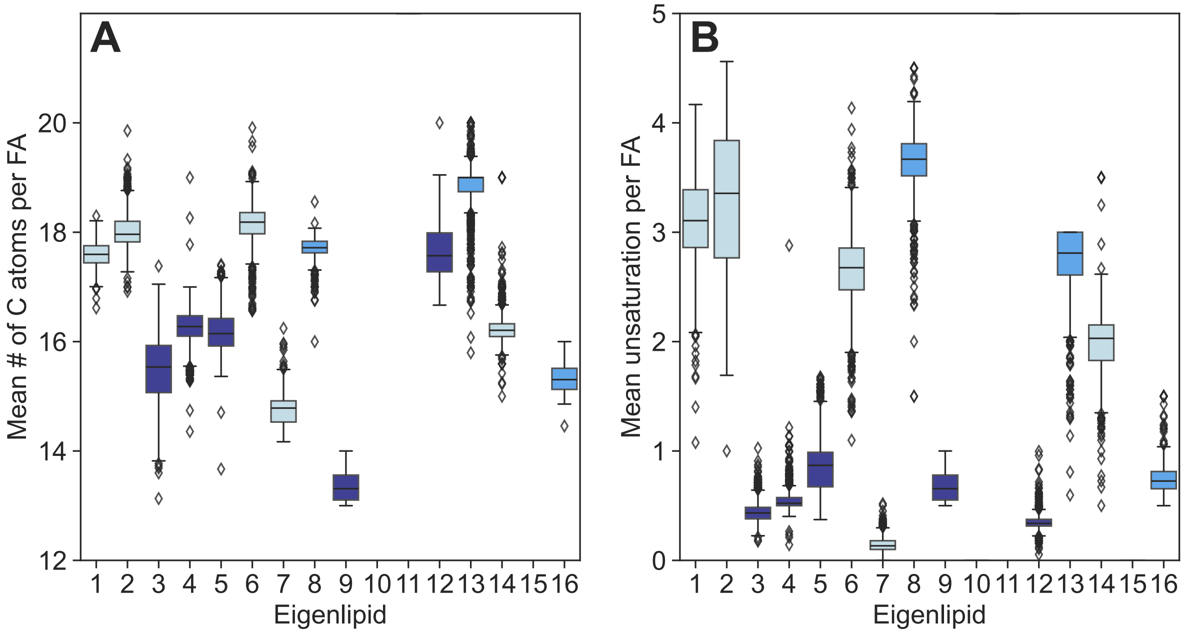


*Figure S2. Distribution of the weighted mean carbon number (A) and unsaturation (B) per fatty acid chains of glycerolipids in each eigenlipid identified* throughout the *sampled* water column *(i.e., both in and below mixed layer). The x-axis denotes the eigenlipid clusters identified by WGCNA. Note that the values refer to the fatty acid moieties excluding the glycerol moieties.*

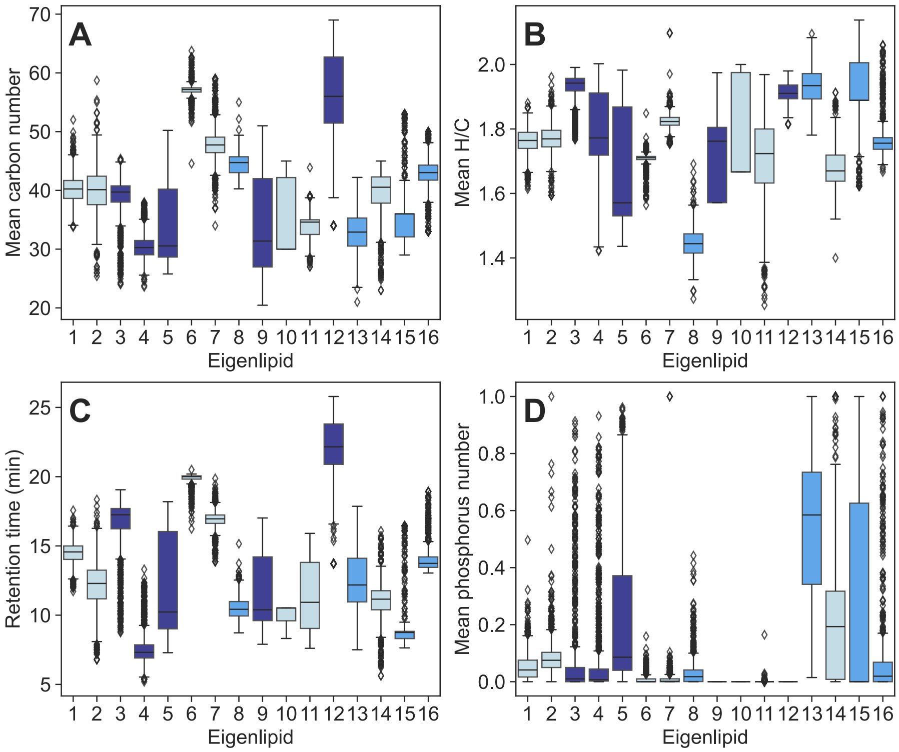


*Figure S3. Molecular properties of Ann. 2 lipids (i.e., excluding the glycerolipids annotated in Holm et al. (2022)) in each eigenlipid cluster identified throughout the sampled water column, include weighted mean carbon number (A), weighted mean hydrogen-to-carbon ratio (B), weighted mean retention time (C), and weighted mean phosphorus number in the formula (D).*

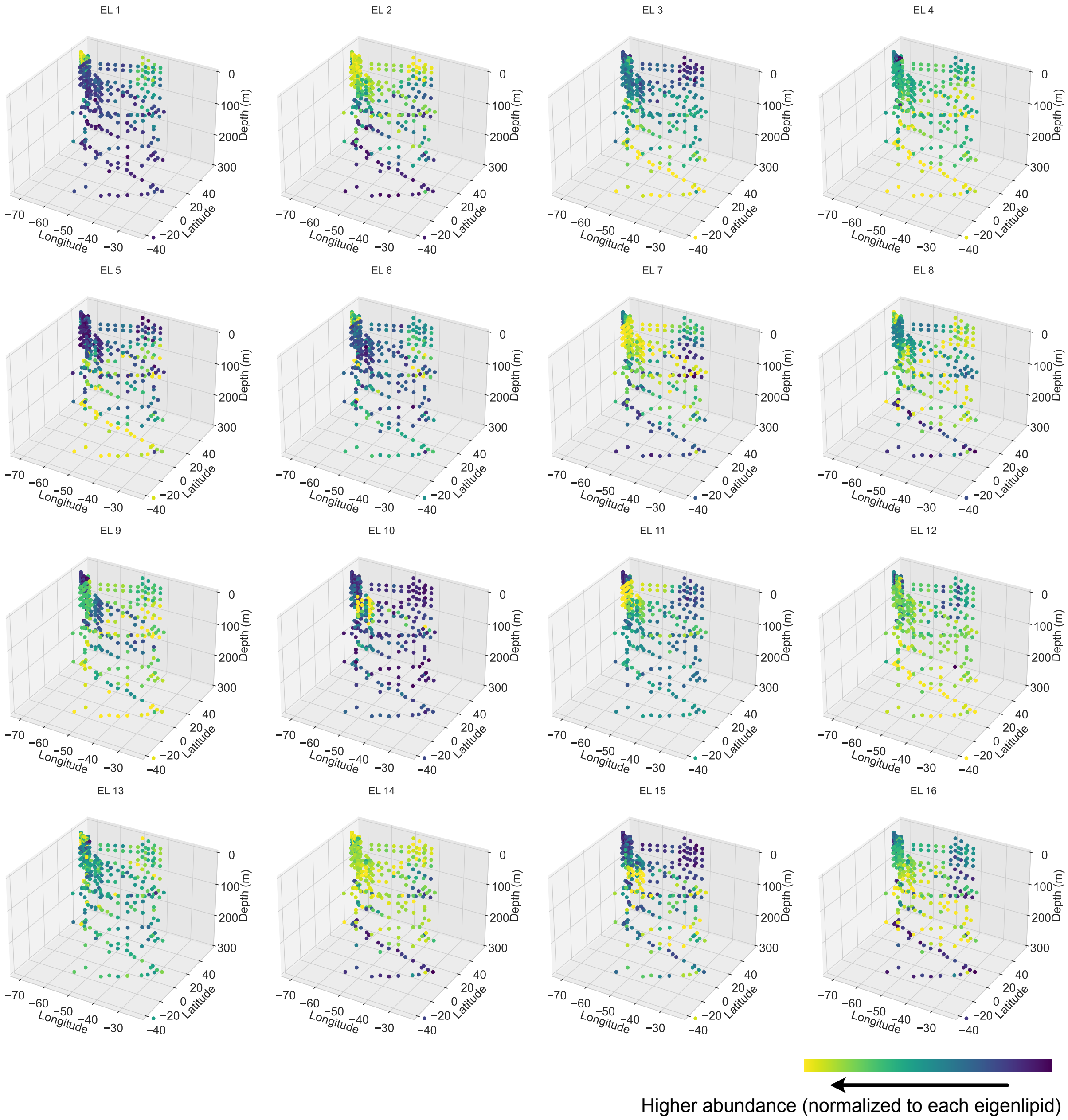


*Figure S4. Spatial distribution of the eigenlipids identified throughout the sampled water column in the Atlantic. The color scale represents the eigenvalue assigned to each eigenlipid cluster.*

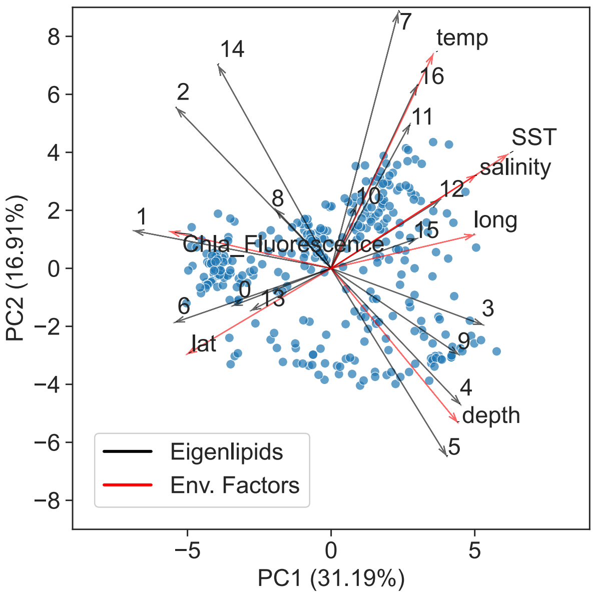


*Figure S5. Principal Component Analysis (PCA) biplot of eigenlipids identified throughout the sampled water column. The biplot displays the first two principal components, PC1 and PC2, accounting for 31.19% and 16.91% of the variance, respectively. Each point represents a sample, and the direction and length of the vectors indicate the contribution and importance of the corresponding variables to the overall variability. The black vectors represent the eigenlipids, and the red vectors represent environmental factors. Note: temp =In-situ temperature, SST = Sea surface temperature, lat = Latitude, long = Longitude*

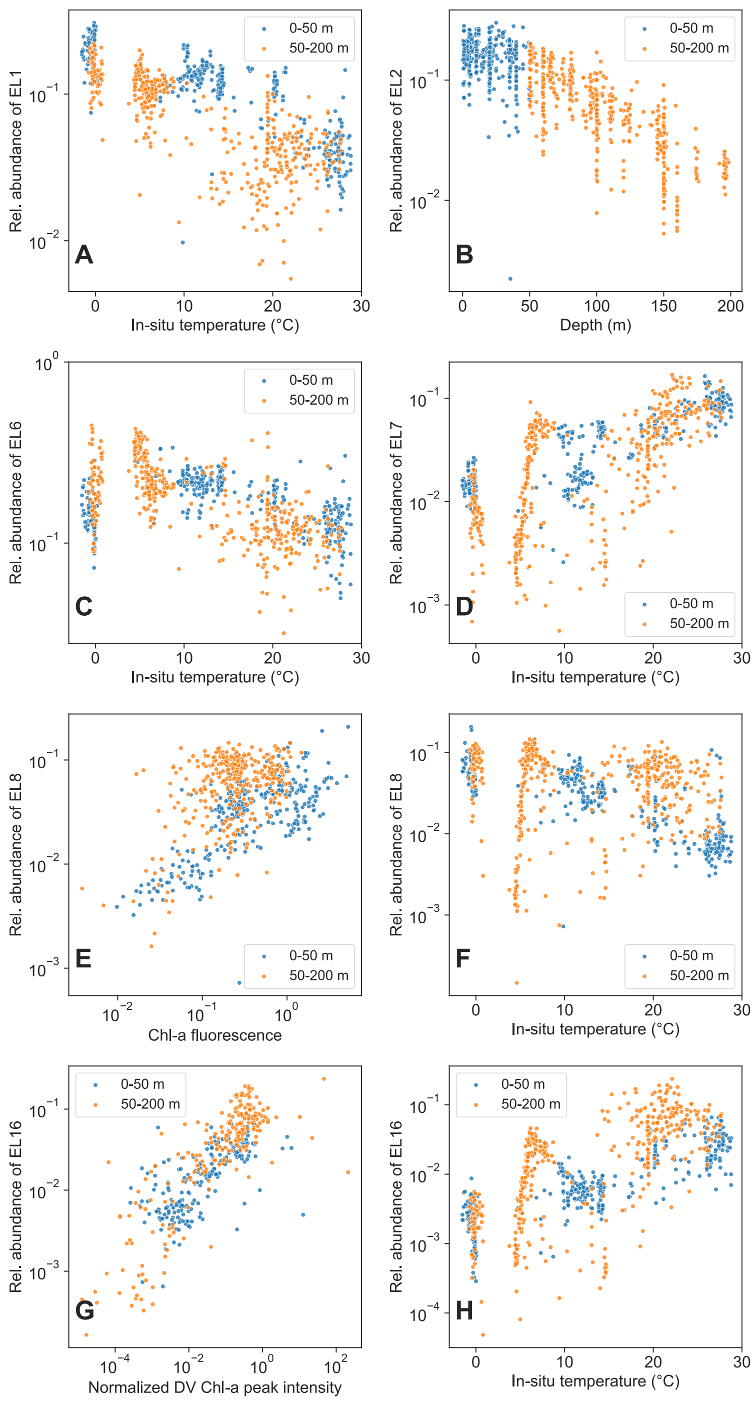


*Figure S6. Influence of environmental forcings on the relative abundance of selected eigenlipids. (A) EL1 vs. In-situ temperature; (B) EL2 vs. Depth; (C) EL6 vs. In-situ temperature; (D) EL7 vs. In-situ temperature; (E) EL8 vs. Chl-a fluorescence; (F) EL8 vs. In-situ temperature; (G) EL16 vs. normalized divinyl chl-a peak intensity (divinyl peak intensity is determined by the peak in extracted ion chromatograms closest at m/z 891.5269, with characteristic fragment ion around m/z 553.21; the peak intensity is then normalized by DNPPE peak intensity; data points from cruise RR1813 and LMG1810 are not shown as the presence of divinyl chl-a cannot be confirmed by MS$^2$ spectra); (H) EL16 vs. In-situ temperature.*

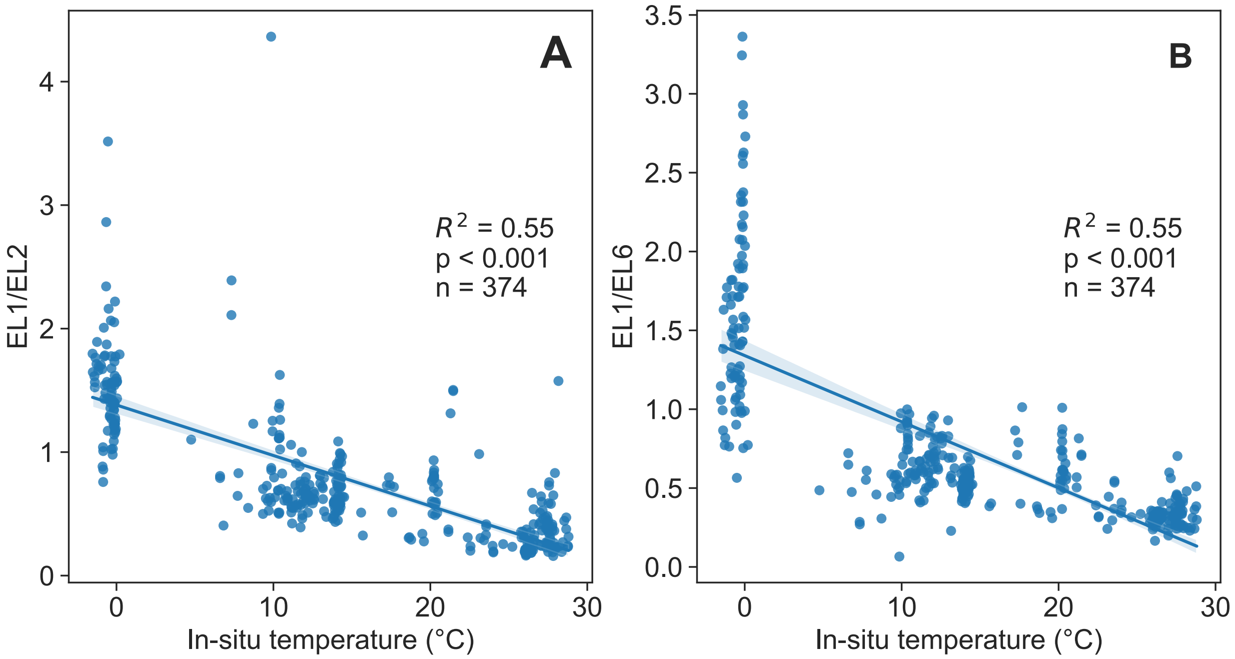


*Figure S7. Stronger response of EL1 lipids to changing temperature than EL2 (A) and EL6 (B), all consisting largely of PUFAs, in the global oceans within a depth range of 50 m. The blue line represents linear fits, and the blue range shows 95% confidence interval of fit. The inset depicts the coefficient of determination for the linear regression fit using ordinary least squares.*

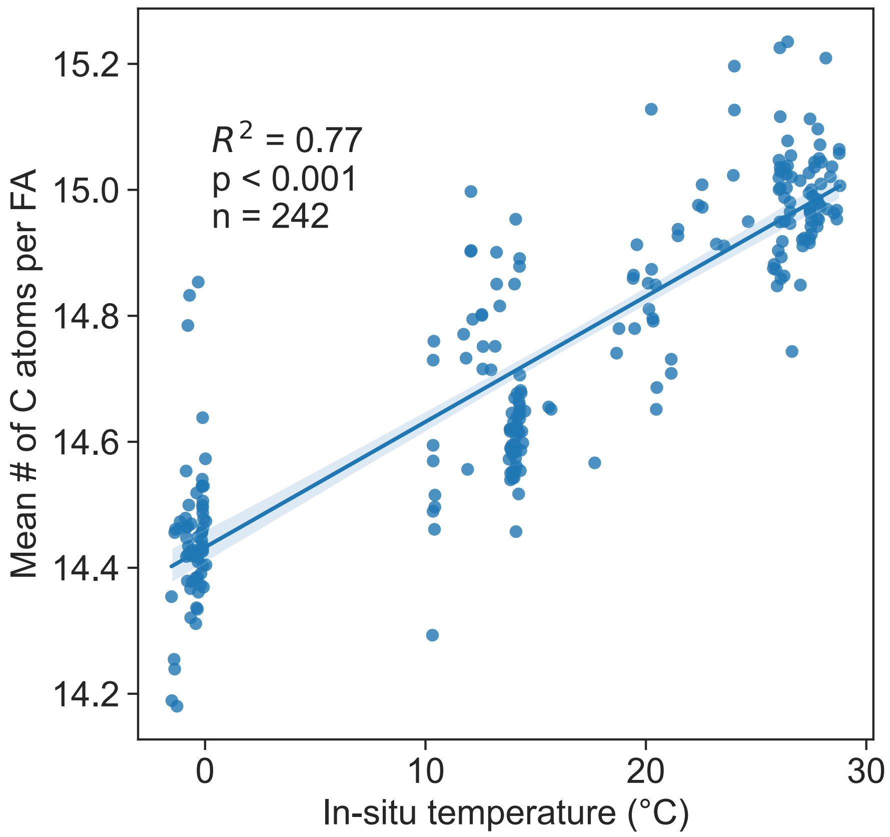


*Figure S8. Trends in fatty acid chain length as a function of temperature across EL7 lipids. The weighted mean carbon number per fatty acid (FA) is plotted against temperature for mixed layer samples. The blue line represents linear fits; the inset depicts the coefficient of determination for the linear regression fit using ordinary least squares.*

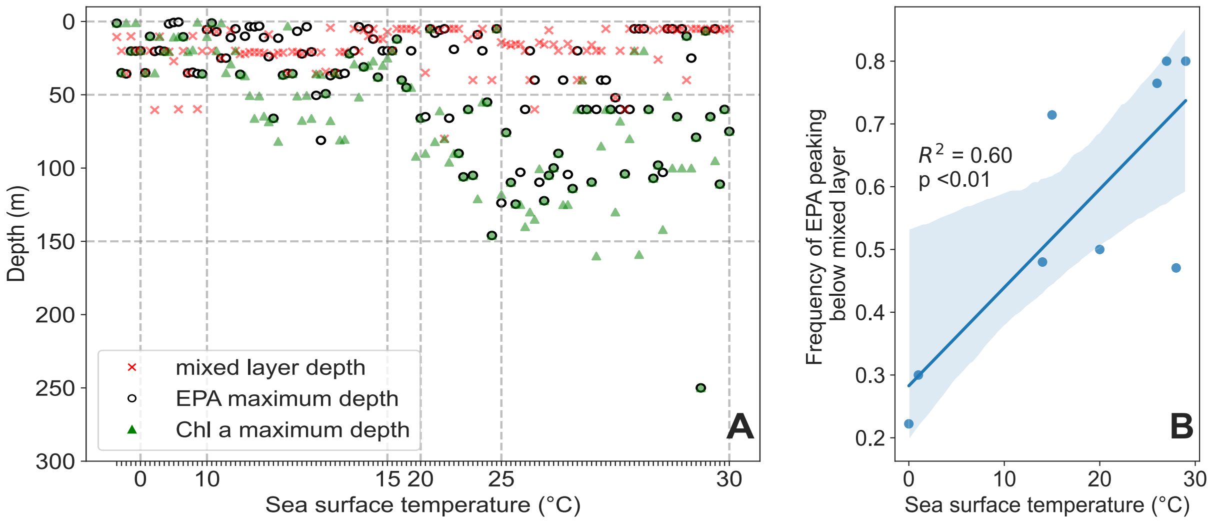


*Figure S9. EPA peaks more frequently below mixed layer near chrlorophyll a maximum in warmer temperature. A: Comparison of mixed layer depth and EPA maximum depth across global oceans. The x-axis represents individual sampled water columns, which are sorted according to sea surface temperature. Open circles denote the depth at which the concentration of EPA-containing species peak in each water column, 'x' marks indicate the mixed layer depth in each water column, and the green triangles denote the depth at which the concentration of chlorophyll-a peaks in each water column. B: Increasing frequency of EPA peak below mixed layer with increasing sea surface temperature. Only the sea surface temperature covered by more than 3 sampling sites is included. The blue line represents linear fits; the inset depicts the coefficient of determination for the linear regression fit using ordinary least squares and the p-value from the F-test (n=9).*

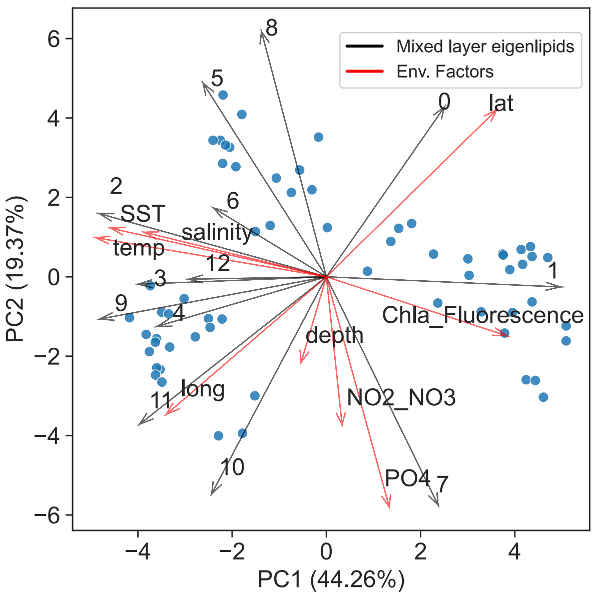


*Figure S10. Principal Component Analysis (PCA) biplot of mixed layer eigenlipids. The biplot displays the first two principal components, PC1 and PC2, accounting for 44.26% and 19.37% of the variance, respectively. Each point represents a sample taken from the Atlantic Ocean, and the direction and length of the vectors indicate the contribution and importance of the corresponding variables to the overall variability. The black and red vectors represent eigenlipids and environmental factors, respectively. Note: temp = In-situ temperature, SST = Sea surface temperature, lat = Latitude, long = Longitude.*

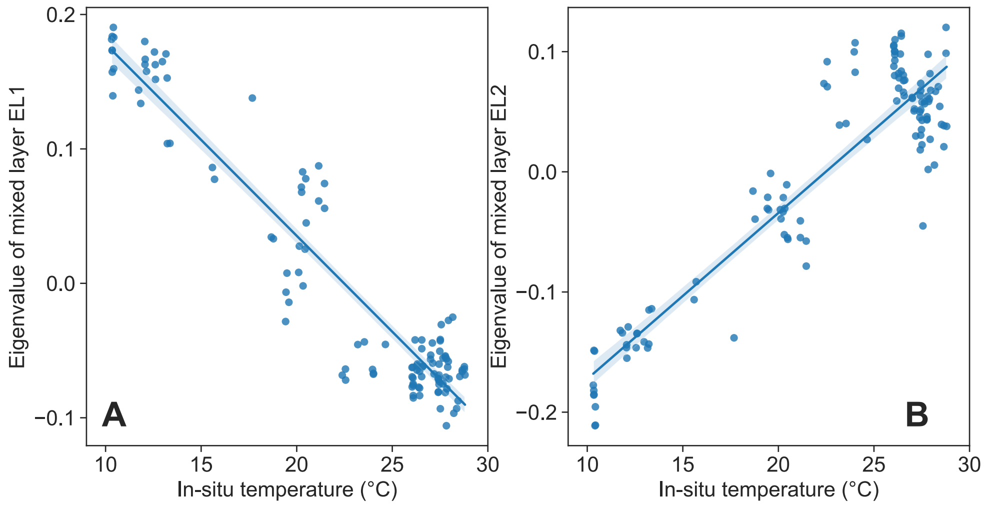


*Figure S11. Correlation between in-situ temperature and mixed layer eigenlipid 1 (A) and 2 (B). Y-axis denotes the eigenvalue of the relative abundance of corresponding eigenlipid cluster.*

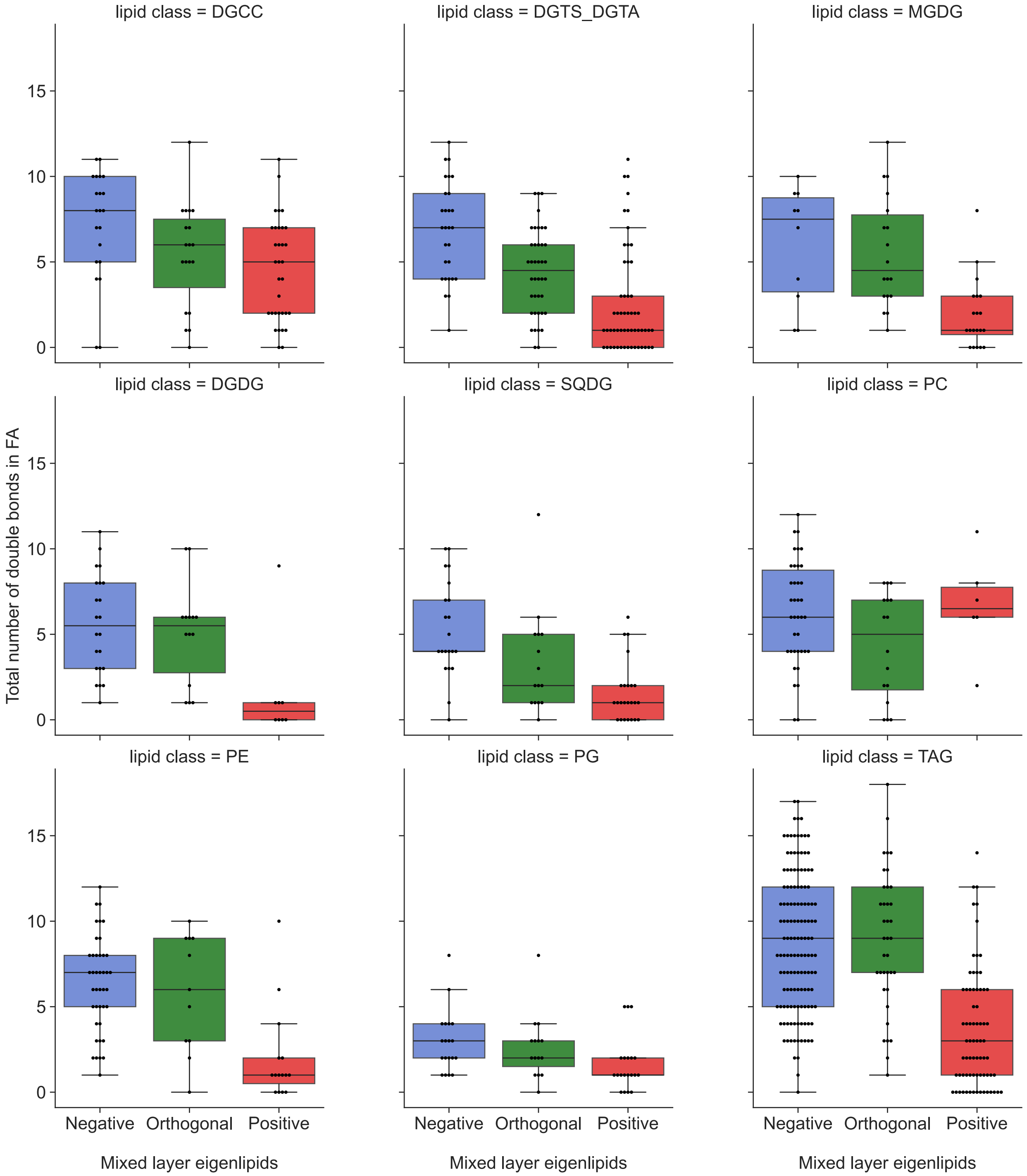


*Figure S12. Distribution of the total number of double bonds in the fatty acid chains of glycerolipids in each mixed layer eigenlipid. The x-axis represents the eigenlipid clusters identified through the WGCNA analysis: 'Negative' represents the eigenlipids that are negatively correlated with temperature, 'Orthogonal' represents the eigenlipids without significant correlation with temperature, and 'Positive' represents the eigenlipids which are positively correlated with temperature.*
